# Microstructural Disruption of the Forceps Minor in Schizophrenia: A Potential Clinical Imaging Biomarker Using Translational DTI

**DOI:** 10.64898/2026.01.28.702301

**Authors:** Qiang Li, Godfrey D. Pearlson, Vince D. Calhoun

## Abstract

Schizophrenia is one of the most complex brain disorders, arising from multidimensional pathophysiological processes that span genetic vulnerability, neurotransmitter dysregulation, structural brain damage, and large-scale brain network dysfunction. Large-scale neuroimaging studies have consistently demonstrated the critical role of frontal brain regions in schizophrenia. Despite substantial progress, the precise localization of structural damage within these regions and the neurobiological mechanisms linking such alterations to disease pathology remain poorly understood. In this study, we included a total of 115 subjects from two sites of the B-SNIP dataset, comprising 60 healthy controls and 55 individuals with schizophrenia. We employed diffusion tensor imaging (DTI) to precisely characterize specific structural alterations in the frontal brain regions associated with schizophrenia. Our findings reveal significant microstructural abnormalities in the forceps minor, a major commissural white-matter tract that serves as a critical interhemispheric bridge between the bilateral frontal lobes. Network-level mapping further demonstrates that the forceps minor is closely integrated with large-scale brain networks, particularly the default mode network, and maintains strong structural connectivity with orbitofrontal regions-both of which are known to exhibit dysfunction in schizophrenia. Moreover, converging evidence suggests that the forceps minor plays an important role in the regulation of social behavior, a core domain of impairment in schizophrenia. Collectively, these findings identify the forceps minor as a promising structural imaging biomarker for schizophrenia and provide novel insights into the microstructural mechanisms underlying the disorder.

## Introduction

Schizophrenia (SCZ) is one of the most complex neurological disorders, involving multidimensional pathophysiological processes, particularly characterized by abnormalities in white matter microstructure, which contribute to disruptions in brain connectivity [1–3]. Accumulating evidence indicates that frontal white matter microstructure reflects heritable abnormalities associated with SCZ [4, 5]. Twin and family studies demonstrate higher monozygotic twin concordance of fractional anisotropy (FA) in frontal tracts, including the forceps minor, genu of the corpus callosum, and anterior cingulum, suggesting a genetic contribution to frontal white matter integrity in SCZ [6]. Structural neuroimaging studies further report frontal system impairments in SCZ [7]. Complementary fMRI studies reveal dysconnectivity within frontal networks and the default mode network [8–11], along with altered orbitofrontal cortex (OFC) activation and volume in SCZ [12–14]. Together, these findings indicate convergent structural and functional abnormalities in frontal brain circuits in SCZ.

However, the contribution of frontal lobe structural abnormalities to the pathophysiology of SCZ has not yet been comprehensively characterized. Substantial gaps remain in our understanding of how damage to precise and specific frontal subregions contributes to the pathophysiology of SCZ.

Diffusion Tensor Imaging (DTI) has emerged as a powerful neuroimaging tool for characterizing the microstructural integrity of white matter in the human brain [15, 16]. Although most SCZ research has traditionally relied on functional MRI (fMRI) to investigate abnormal brain activity and connectivity, DTI provides complementary information by directly assessing the structural pathways that support these networks [17, 18]. Subtle disruptions in white matter microstructure, detectable through DTI, may serve as reliable imaging biomarkers for SCZ [19–21]. By elucidating these structural alterations, DTI not only enhances our understanding of the neurobiological underpinnings of SCZ but also holds promise for improving diagnosis and informing targeted clinical interventions.

The primary objective of this study is to localize specific frontal structural alterations associated with SCZ using DTI. By integrating functional connectomic findings with fMRI measures and behavioral assessments derived from small-animal experiments, this study aims to elucidate the underlying neurobiological mechanisms of SCZ at the macroscale level. Ultimately, this work seeks to advance mechanistic understanding of SCZ and to inform the longterm development of more targeted and effective clinical interventions.

## Materials and Methods

### Participants

Data included in this study were collected from 115 participants from two sites (Hartford, Baltimore) of the Bipolar-Schizophrenia Network on Intermediate Phenotypes (B-SNIP) consortium [22, 23]: 60 CNs, 55 subjects with SCZ (Table. I). All participants provided written informed consent statements approved by review boards of Hartford Hospital/Yale University and the University of Maryland/Johns Hopkins University.

**TABLE I:** Participant demographics.

| Measure | CN | SCZ |
| --- | --- | --- |
| **N** | 60 | 55 |
| **N Hartford** | 23 | 26 |
| **N Baltimore** | 37 | 29 |
| **Mean Age/SD** | 37.41/9.05 | 32.93/8.03 |
| **Male/Female** | 28/32 | 38/17 |
| Note: Standard deviation (SD) |  |  |

The consensus clinical diagnoses for SCZ were made by trained clinical raters and senior psychiatric diagnosticians using clinical data and the Structured Clinical Interview for DSM-IV (SCID). CNs were screened and found to be free of known neurological illnesses and not actively using illicit substances, as confirmed by negative urine toxicology screens.

### Diffusion Weighted Imaging Acquisition

Diffusion-weighted imaging (DWI) data were acquired using 3T MRI scanners. Scanners located at Hartford (Siemens Allegra) and Baltimore (Siemens Triotim) employed similar scanning sequences. Both sites utilized single-shot spin-echo planar imaging with a twice-refocused balanced echo sequence to minimize eddy current distortions. Specifically, the scanning parameters for the Hartford site included a repetition time (TR) of 6300 ms, an echo time (TE) of 85 ms, a field of view (FOV) of 220 mm, a b-value of 1000 s/mm^2^ across 32 diffusion directions, 45 contiguous slices, three imaging series, and a voxel size of 1.7 × 1.7 × 3 mm. For the Baltimore site, the parameters were a TR of 6700 ms, a TE of 92 ms, a FOV of 230 mm, a b-value of 1000 s/mm^2^ across 30 diffusion directions, 48 contiguous slices, two imaging series, and a voxel size of 1.8 × 1.8 × 3 mm.

### Preprocessing and Postprocessing

DWI were carefully preprocessed to ensure data quality and anatomical consistency across subjects. First, data were acquired across two sites with different scanners, so we applied a harmonization technique [24] that removes scanner-related effects while preserving inter-subject biological variability and group differences, working directly on the raw diffusion MRI signal in a model-independent manner. Then, the preprocessing included correction for eddy current distortions and subject motion, followed by alignment of the diffusion volumes to the anatomical reference. All data were spatially normalized to the standard MNI-EPI template to provide a common coordinate space across participants, and the diffusion gradient vectors were adjusted to account for these spatial transformations, ensuring accurate representation of fiber orientations. After preprocessing, the diffusion tensor was fitted at each voxel using a trilinear interpolation approach, which estimates the tensor from the weighted diffusion signal.

From the fitted diffusion tensor, voxel-wise maps of fractional anisotropy (FA), mean diffusivity (MD), axial diffusivity (AD), and radial diffusivity (RD) can be computed, providing quantitative measures of microstructural properties. Disruptions in the myelination process, such as damage to the myelin sheath, can alter these DTI metrics, reflecting impaired axonal integrity or incomplete myelination (Fig.1). Such changes can disrupt normal brain function and connectivity, making DTI a valuable tool for assessing myelination levels and monitoring the progression of SCZ. See the ***Supplement*** for details regarding data quality control and diffusion metric calculations.

**FIG. 1:**
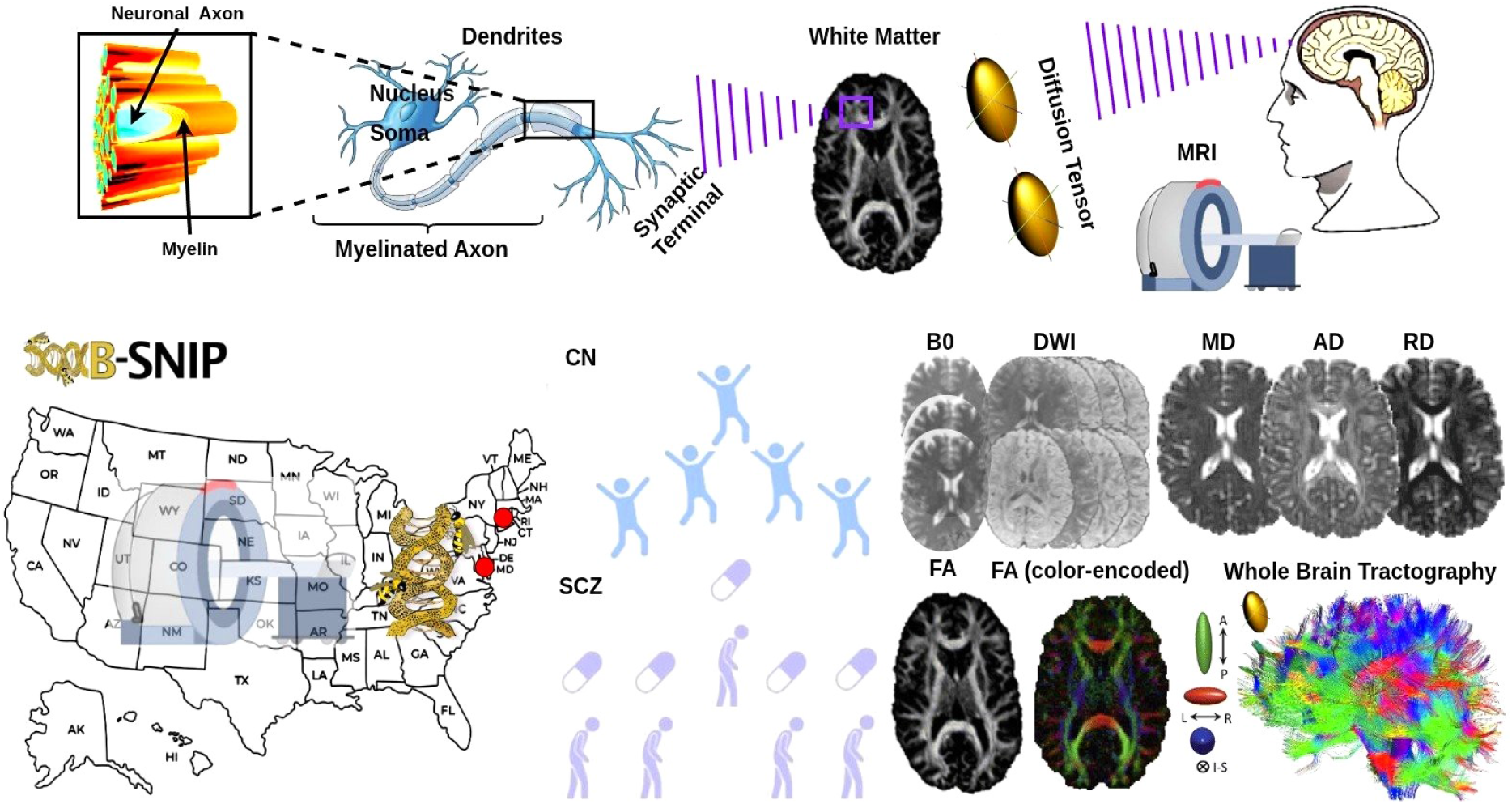
Study overview. DTI helps visualize and quantify the integrity of white matter by tracking water movement along myelinated axons. Disruptions in myelination, such as damage to the myelin sheath, can alter DTI metrics, reflecting impaired axonal integrity and potentially disrupting brain function and connectivity. The B-SNIP dataset, collected from Hartford (Connecticut) and Baltimore (Maryland), was used in this study. Healthy controls (CN) and SCZ were selected, and corresponding *B*_0_ and diffusion weighted imaging data were acquired for both cohorts. After standardized preprocessing and follow up analyses, MD, AD, RD, and FA maps (color-encoded fibers: blue = superior-inferior, green = anterior-posterior, red = left-right) were generated. Whole brain tractography was subsequently performed, and quantitative analyses were applied to characterize and differentiate microstructural abnormalities between the CN and SCZ groups.

### Statistical Analyses

To assess differences between SCZ and CN, a two sample t-test was conducted for each of the 20 fiber tracts based on the JHU white matter tractography template [25] (see ***Supplement*** for details), comparing values of FA, MD, RD, and AD at each point along the tracts. The results, including t-statistics and p-values, were computed to identify significant differences between the groups. These statistical outputs were incorporated into the tract profiles, allowing us to identify specific points along the fiber tracts where significant differences in FA, MD, RD, or AD existed between SCZ and CN. The p-values indicated the statistical significance of these differences, while the t-statistics provided insight into the magnitude and direction of the differences. This comprehensive analysis allowed us to better understand how each of these diffusion metrics varied across the different fiber tracts, highlighting areas with significant group differences.

## Results

### Forceps Minor FA Abnormalization in SCZ

In our study, we observed FA abnormalities in the forceps minor in patients with SCZ, reflecting altered microstructural integrity of interhemispheric frontal white matter connections (Fig.2). FA was regionally heterogeneous, with decreases in some areas affecting the overall fiber integrity; nevertheless, the overall FA in the forceps minor differed significantly between SCZ patients and CN (*p <* 0.05, FDR-corrected *p* = 0.01) compared to other fiber tracts (Fig.3), indicating disrupted structural connectivity within frontal networks. Reduced FA generally reflects impaired axonal organization, decreased myelination, or altered fiber coherence. These findings support the frontal dysconnectivity hypothesis of SCZ, which proposes that disrupted communication among distributed brain regions underlies core cognitive and clinical symptoms of the disorder.

**FIG. 2:**
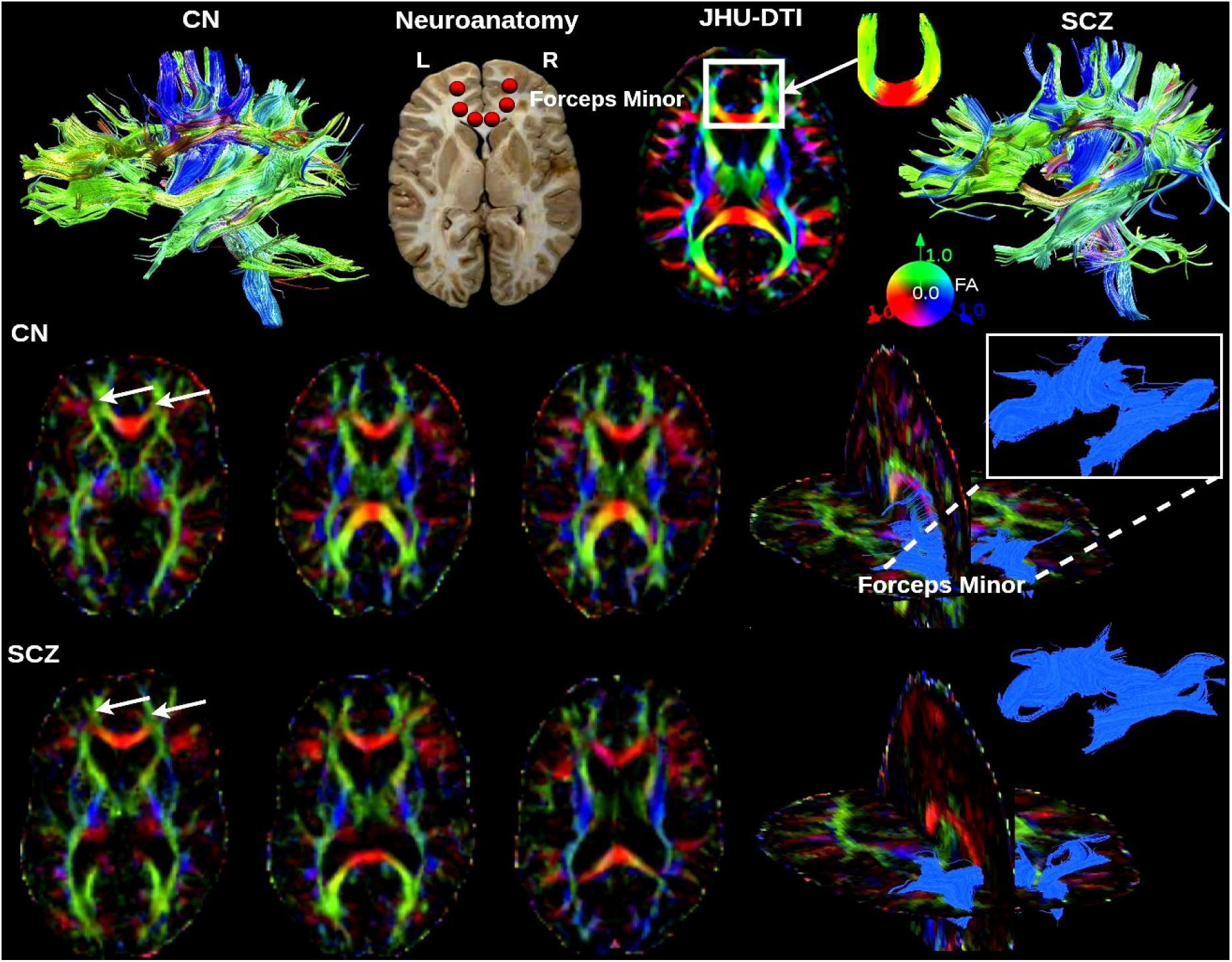
Whole brain fiber tracts in CN and SCZ. Whole brain fiber tracts for both CN and individuals with SCZ are presented. To optimize rendering time, only 6,000 fibers were included in the display. The neuroanatomy of the forceps minor is labeled for comparison with its corresponding regions on the FA maps, which are shown on the JHU-DTI FA template. Additionally, FA maps for both the CN and SCZ groups are shown as axial slices and 3D meshes, with the forceps minor highlighted by arrows and its segmented portion indicated.

**FIG. 3:**
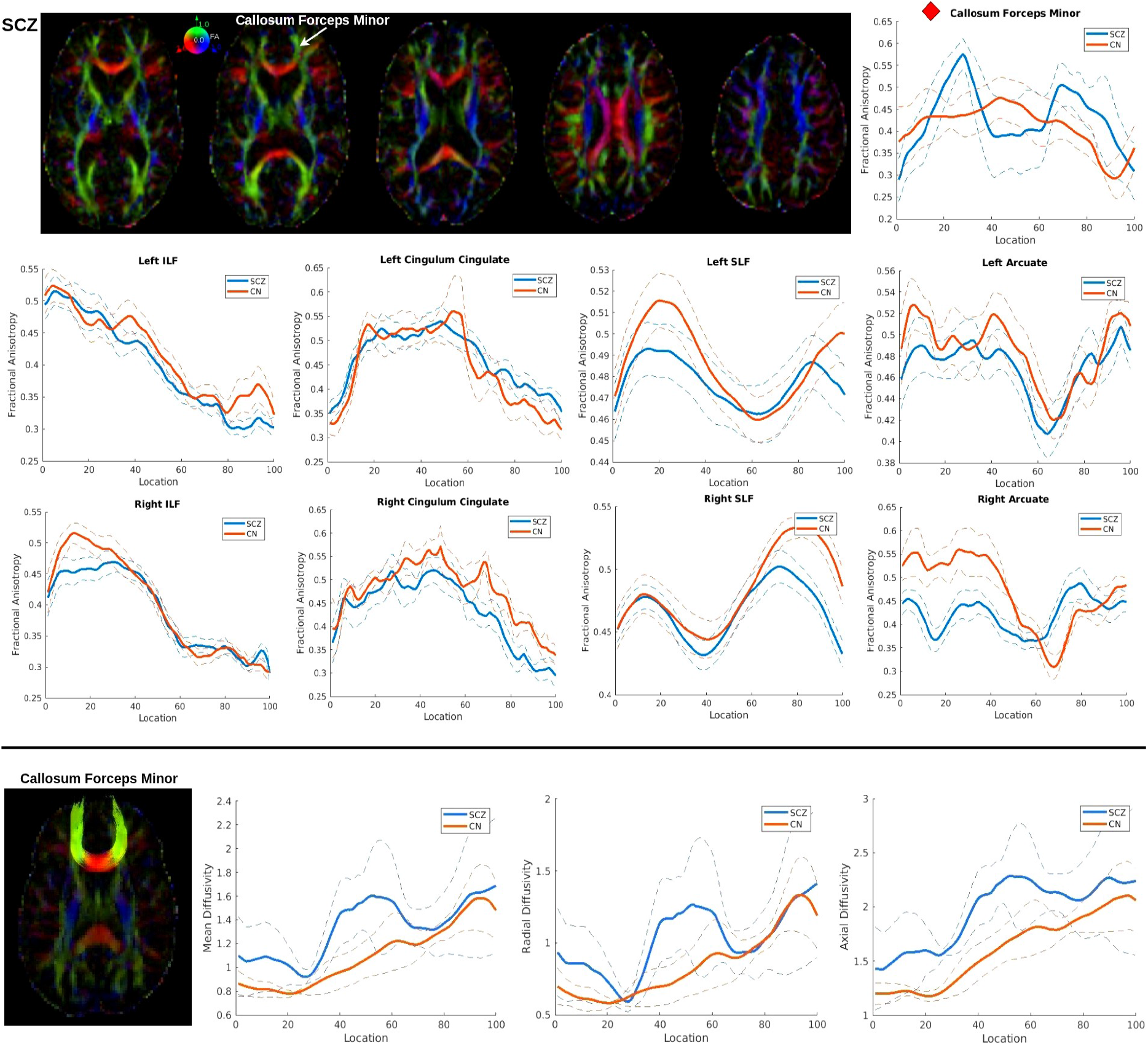
Group differences in pointwise white matter fiber tract metrics between SCZ and CN. The forceps minor of the corpus callosum, a white matter tract connecting the frontal regions of the left and right hemispheres, showed significant differences in FA between SCZ and CN (*p <* 0.05, FDR-corrected), compared with other white matter tracts. Additionally, from left to right in the second row, the plots display MD, RD, and AD values along the forceps minor, highlighting segments that show significant group differences between SCZ and CN (*p <* 0.05, FDR-corrected). The blue line represents the SCZ group, and the orange line represents the CN group; solid lines indicate group means, and dotted lines represent standard deviations.

### Forceps Minor MD, RD, and AD Abnormalization in SCZ

In addition to FA abnormalities, we observed alterations in MD, RD, and AD within the forceps minor in SCZ (Fig.3), providing further insight into the underlying microstructural pathology. Increased MD in this tract indicates reduced overall tissue integrity, potentially reflecting neurodevelopmental disturbances or progressive white matter degeneration. Elevated RD is commonly interpreted as evidence of myelin disruption or dysmyelination, whereas alterations in AD may reflect axonal injury or changes in axonal caliber. Collectively, these diffusivity measures complement FA findings and underscore the multifaceted nature of white matter abnormalities in SCZ.

### Functional Relevance of the Forceps Minor

The forceps minor has multiple functional roles (Fig.4), playing a critical part in interhemispheric communication between prefrontal regions, including the orbitofrontal cortex (OFC), which has been reported as abnormal in SCZ [12–14]. From a network perspective, structural abnormalities in the forceps minor may impair the efficiency and integration of key large-scale networks, particularly the default mode network (DMN), specifically within the medial prefrontal cortex (mPFC) [26].

**FIG. 4:**
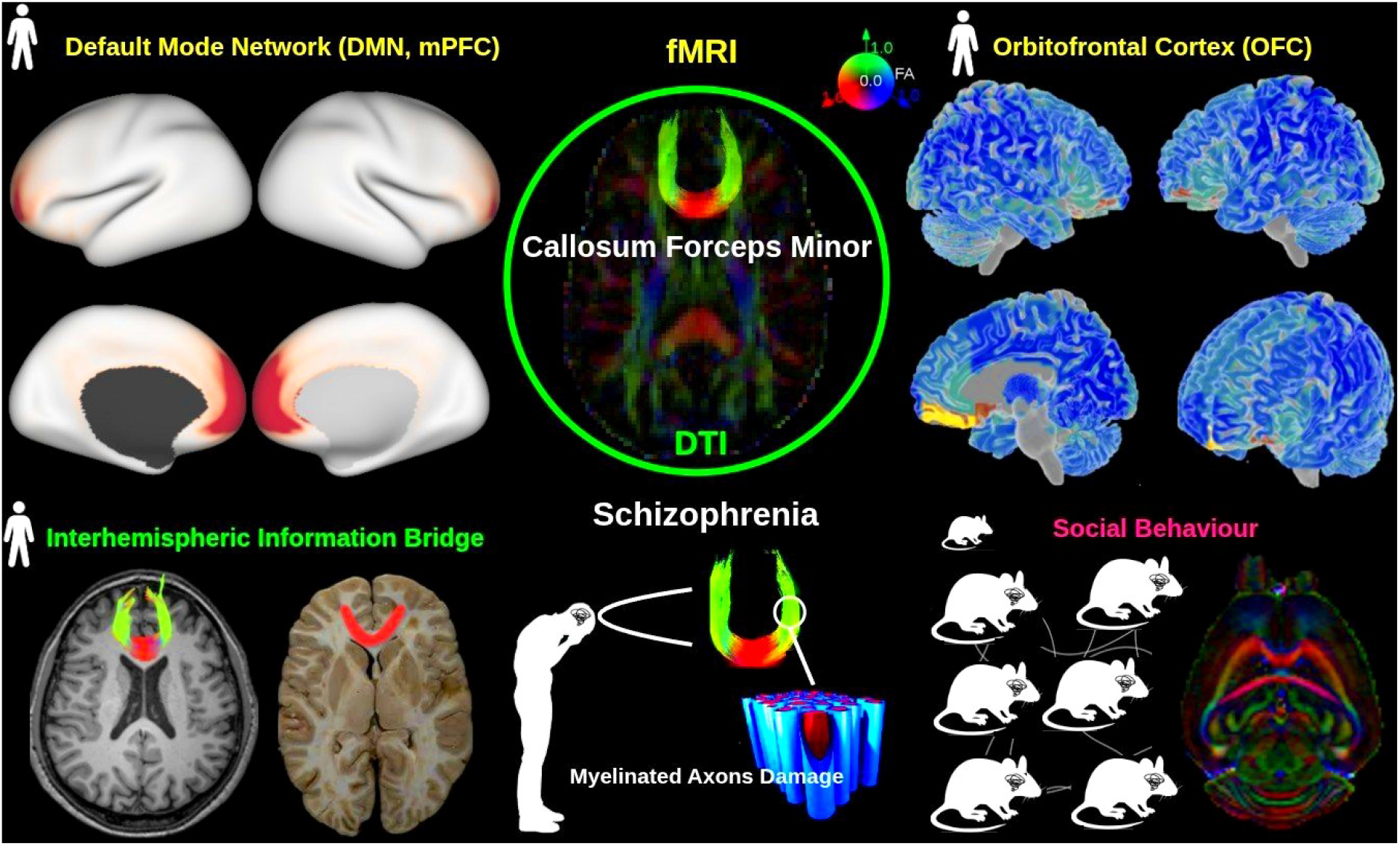
Connectivity and functional relevance of the forceps minor. The forceps minor is primarily associated with the DMN, particularly the mPFC, and shows prominent connections with the OFC. By linking homologous frontal regions across hemispheres, it functions as a key interhemispheric pathway supporting frontal network integration. Consistent with these roles, evidence from small-animal studies suggests that the forceps minor is involved in social behavior. Accordingly, microstructural abnormalities of the forceps minor contribute to disrupted frontal connectivity and the pathophysiology of schizophrenia.

Disruptions in the structure of the forceps minor can lead to altered connectivity within the DMN, which in turn may contribute to the functional connectivity abnormalities observed in SCZ [27, 28]. This creates a cascade from structural deficits to network dysfunction, ultimately contributing to the clinical symptoms of the disorder.

Functionally, the forceps minor is implicated in higherorder cognitive processes, including social behavior. Preclinical studies [29] suggest that demyelination of the forceps minor is associated with increased anxiety and decreased social interaction in animal models, whereas remyelination can restore normal social behaviors.

***Outstanding Questions***

What are the specific cellular and molecular mechanisms that contribute to the white matter changes observed in SCZ? While DTI provides valuable insights into structural connectivity, it remains unclear whether these changes are primarily due to myelin loss, axonal injury, or other forms of damage. Understanding the precise underlying mechanisms is crucial for interpreting DTI results and advancing SCZ research.

How do white matter changes in SCZ, particularly in tracts like the forceps minor, correlate with specific cognitive and behavioral impairments Is there a direct link between disruptions in the forceps minor tract and the cognitive symptoms or behavioral issues observed in SCZ patients?

What are the interactions between gray and white matter abnormalities in SCZ, and how do they together contribute to the pathophysiology of the disorder? Could integrating findings from both gray and white matter changes offer a more holistic view of SCZ, and how can imaging techniques be refined to better capture these interrelationships?

Are there novel therapeutic approaches that could specifically target white matter abnormalities in SCZ, potentially reversing or halting the progression of these changes? Could targeting specific molecular pathways involved in white matter damage (e.g., myelin repair or axonal regeneration) open new therapeutic avenues for improving cognitive and functional outcomes in SCZ patients?

These findings highlight the importance of the forceps minor in mediating both cognitive and social functions, and suggest that its structural integrity is critical for the proper functioning of frontal networks affected in SCZ.

## Discussion

In the current study, we investigated the microstructure of white matter in individuals with SCZ using DTI. Specifically, we aimed to identify disruptions in fiber tracts that are associated with this disorder. By examining diffusion metrics such as FA, MD, RD, and AD, we observed significant alterations in white matter integrity. Notably, we identified damage to the forceps minor, a crucial white matter tract involved in interhemispheric communication. These findings suggest that forceps minor damage may serve as a potential imaging biomarker for SCZ, offering valuable insights for diagnosis and understanding the pathophysiology of the disorder.

The forceps minor is a major interhemispheric white matter tract connecting the bilateral medial and orbitofrontal regions of the frontal lobe [30]. These regions include key nodes of the DMN, particularly the mPFC [26]. Disruption of the forceps minor may therefore impair interhemispheric integration of prefrontal areas, leading to altered structural support for DMN connectivity. The OFC, which is anatomically and functionally connected with the mPFC, plays a critical role in emotional regulation [31, 32], and decision-making [33]. Moreover, demyelination of the forceps minor has been shown to impair social behavior, leading to a decrease in social interactions [29]. However, subsequent remyelination appears to facilitate the re-establishment of social behavior, providing evidence that white matter integrity in this tract plays a critical role in the modulation of social cognitive functions [29]. Therefore, structural damage to the forceps minor may compromise communication between bilateral mPFC and OFC regions, potentially contributing to DMN dysfunction and prefrontal network abnormalities observed in SCZ.

One limitation of this study is the challenge of interpreting DTI results in SCZ, particularly when it comes to understanding white matter changes. While DTI offers valuable insights into the brain’s structural connectivity, it does not directly reveal the underlying causes of damage, e.g., whether the changes are due to myelin loss or axonal injury [34]. This can make it harder to draw precise conclusions, especially given the complex nature of SCZ, which involves both gray and white matter abnormalities. To gain a clearer understanding, future research could benefit from combining DTI with other imaging techniques, such as myelin-specific imaging or histological analysis, which could provide more detailed insights into the structural changes that contribute to cognitive and behavioral issues in SCZ.

For future studies, it also would be valuable to incorporate small-animal mouse models to validate the findings of white matter changes observed in SCZ. For example, using genetic SCZ mouse models could help investigate whether damage to the forceps minor occurs in a similar manner to what we observed in human subjects. DTI could be employed in these models to confirm the specific disruptions in the forceps minor tract and further explore the underlying mechanisms of these changes. This approach would allow for more controlled investigations of how forceps minor damage contributes to cognitive and behavioral symptoms in SCZ, providing a solid foundation for understanding the pathophysiology of the disorder.

## Conclusion

This study has identified the forceps minor as a structural biomarker, offering valuable insights into the neurobiological mechanisms underlying SCZ through DTI. Damage to the microstructure of frontal brain white matter, particularly within the forceps minor, was observed in individuals with SCZ. Furthermore, the forceps minor has been shown to connect with functional brain networks and regions, playing a role in regulating social behavior. These findings suggest that the forceps minor could serve as a potential early clinical structural biomarker and sign for SCZ diagnosis.

Although several fundamental questions remain (see ***Outstanding Questions*** ), we have identified the forceps minor as a potential imaging biomarker for clinical applications and demonstrated the utility of translational DTI for studying SCZ.

## Acknowledgments

This work was supported by the National Science Foundation grant 2112455, and the National Institutes of Health grants R01MH123610 and R01MH119251.

## Author Contributions

Q. Li., VD. Calhoun.: Conceptualization, Investigation, Software, Writing - Review & editing. Q. Li., GD. Pearlson., VD. Calhoun.: Writing - Review & editing. VD. Calhoun.: Funding Acquisition.

